# Mutation of the ATPase domain of MutS homolog-5 (MSH5) reveals a requirement for a functional MutSγ complex for all crossovers in mammalian meiosis

**DOI:** 10.1101/546010

**Authors:** Carolyn R. Milano, J. Kim Holloway, Yongwei Zhang, Bo Jin, Aviv Bergman, Winfried Edelmann, Paula E. Cohen

## Abstract

During meiosis, induction of DNA double strand breaks (DSB) leads to recombination between homologous chromosomes, resulting in crossovers (CO) and non-crossovers (NCO). Only 10% DSBs resolve as COs, mostly through a class I pathway dependent on MutSγ (MSH4/ MSH5). Class II CO events represent a minor proportion of the total CO count and also arise from DSBs, but are not thought to involve MutSγ. However, loading of MutSγ occurs very early in prophase I at a frequency that far exceeds the final number of class I COs found in late prophase I. Moreover, loss of MutSγ in mouse results in apoptosis before CO formation, preventing analysis of its CO function. We generated a mutation in the ATP binding domain of *Msh5* (*Msh5^GA^*). While this mutation was not expected to affect MutSγ complex formation, MutSγ foci do not accumulate during prophase I. Nevertheless, while some spermatocytes from *Msh5^-/-^* animals progress into pachynema, most spermatocytes from *Msh5^GA/GA^* mice progress to late pachynema and beyond. Some spermatocytes from *Msh5^GA/GA^* mice complete prophase I entirely, allowing for the first time an assessment of MSH5 function in CO formation. At pachynema, *Msh5^GA/GA^* spermatocytes show persistent DSBs, incomplete homolog pairing, and fail to accumulate MutLγ (MLH1/MLH3). Unexpectedly, *Msh5^GA/GA^* diakinesis-staged spermatocytes have no chiasmata at all from any CO pathway, indicating that a functional MutSγ complex in early prophase I is a pre-requisite for all COs.

**ARTICLE SUMMARY:** MSH4/MSH5 are critical components of the class I crossover (CO) machinery, which is responsible for >90% of the COs that arise in mammalian meiosis. We generated a point mutation in the ATP binding motif of *Msh5*, and found that mutant spermatocytes lose all COs, not just those arising from the class I pathway.

## INTRODUCTION

MSH5 (MutS homolog 5) belongs to the DNA mismatch repair (MMR) family of proteins that perform multiple DNA repair activities, most prominently the correction of mispaired bases that result from erroneous DNA replication (Modrich and Lahue 1996). Like other family members, MSH5 acts with a MutS homolog partner, specifically with MSH4, to form the MutSγ heterodimer (Bocker *et al.* 1999). Unlike other MutS heterodimers, MutSγ does not participate in mismatch correction in somatic cells, but instead acts exclusively during meiotic prophase I in budding yeast (Pochart *et al.* 1997), mice (Edelmann *et al.* 1999; de Vries *et al.* 1999; Kneitz *et al.* 2000; Santucci-Darmanin and Paquis-Flucklinger 2003), humans (Bocker *et al.* 1999), plants (Higgins, Vignard, *et al.* 2008), and worms (Zalevsky *et al.* 1999). Importantly, loss of either MutSγ subunit results in infertility among mammals, including humans and mice (Edelmann *et al.* 1999; de Vries *et al.* 1999; Kneitz *et al.* 2000; Carlosama *et al.* 2017).

Prophase I is the defining stage of meiosis, encompassing the unique events that give rise to pairing and equal segregation of homologous chromosomes at the first meiotic division. In early prophase I homologous chromosomes undergo a physical tethering process known as *synapsis.* Synapsis is mediated by the proteinaceous structure called the Synaptonemal Complex (SC) whose status defines the sub-stages of prophase I: leptonema, zygonema, pachynema, diplonema, and diakinesis. Synapsis is dependent on, and facilitated by, homologous recombination, which is triggered by the formation of DNA double strand breaks (DSBs) by the transesterase, SPO11 and its co-factors (Keeney *et al.* 1997; Baudat *et al.* 2000; Romanienko and Camerini-Otero 2000; Keeney 2008). DSBs are processed sequentially through various common repair intermediates including the resection of one end of the DSB and its invasion into an opposing homolog to induce a displacement loop (D-loop) in the recipient molecule (Hunter 2015). This intermediate may then be resolved through multiple distinct, yet overlapping, pathways that result in either a crossover (CO) or a non-crossover (NCO)(Gray and Cohen 2016). While CO repair is essential for the maintenance of homolog interactions until the first meiotic divisions, the large number of DSBs that form, whether resulting in COs or NCOs, are thought to be essential for enabling homolog recognition and pairing in early prophase I. In mouse, the majority (approximately 90%) of the 250+ DSBs that form are processed to become NCOs (Cole *et al.* 2014), and these are thought to arise largely out of the Synthesis Dependent Strand Annealing (SDSA) pathway. In yeast, and probably also in the mouse, SDSA utilizes early D-loop intermediates and the ensuing NCOs arise at temporally earlier time points than do the CO repair products (Allers and Lichten 2001; Baudat and de Massy 2007; Jessop and Lichten 2008; Kaur *et al.* 2015).

DSB-induced COs result in reciprocal exchange of DNA between maternal and paternal homologs, and give rise to the chiasmata that ensure the maintenance of homologous chromosome interactions until the first meiotic division. In mouse, only 20-30 (10%) become repaired as COs, via two distinct pathways that have been characterized in budding yeast, plants, and mouse (de los Santos *et al.* 2003; Argueso *et al.* 2004; Mercier *et al.* 2005; Higgins, Buckling, *et al.* 2008; Holloway *et al.* 2008).

The first is the major Class I pathway that involves the ZMM group of proteins, of which the MutSγ constituents are members, along with Zip1-4, Mer3, and Spo16 (Lynn *et al.* 2007). This pathway involves single, end invasion (SEI), second end capture of the other side of the DSB, and subsequent formation of a double Holliday Junction (dHJ) that must then be resolved via the action of resolvases which cleave the dHJs to release the two recombined homologous chromosomes. The class I pathway is dependent on a second MMR heterodimer consisting of MLH1 and MLH3 (collectively known as MutLγ), which has been proposed to be at least one resolvase that is active in meiosis (Edelmann *et al.* 1996; Hunter and Borts 1997; Wang *et al.* 1999; Lipkin *et al.* 2002; Svetlanov *et al.* 2008; Nishant *et al.* 2008). According to the canonical mechanism of MMR recruitment, specific MutS heterodimers recruit specific MutL heterodimers to the site of the DNA lesion, a model that fits with the temporally distinct localization of MutSγ and MutLγ in mammalian meiosis (Moens *et al.* 2002; Santucci-Darmanin and Paquis-Flucklinger 2003). Indeed, MLH1 and MLH3 localize to meiotic chromosomes cores during pachynema of prophase I, and their localization defines the final cohort of class I CO events (Baker *et al.* 1996; Edelmann *et al.* 1999; Kneitz *et al.* 2000; Lipkin *et al.* 2002; Santucci-Darmanin and Paquis-Flucklinger 2003; Kolas *et al.* 2005).

The second CO pathway, the class II pathway, is dependent on the MUS81-EME1 endonuclease, and in many organisms accounts for only a fraction of the total CO count (Oh *et al.* 2008; Holloway *et al.* 2008). This pathway does not involve canonical dHJ formation but instead may resolve a diverse set of repair intermediates that would not ordinarily be strong substrates for the class I machinery. In mice, the class I CO pathway ultimately accounts for more than 90% of COs and these events are considered to be “interference” dependent, meaning that the placement of one CO prevents nearby COs. The class II pathway accounts for the remaining 10%, and such events are not sensitive to interference (Edelmann *et al.* 1996; Lipkin *et al.* 2002; Lynn *et al.* 2007; Holloway *et al.* 2008).

The role of MLH1 and MLH3 in class I COs has been demonstrated by the fact that *Mlh1^-/-^* and *Mlh3^-/-^* mice show mostly univalent chromosomes, with a 90% reduction in the number of chiasmata (Edelmann *et al.* 1996; Lipkin *et al.* 2002). By contrast, loss of *Msh4* or *Msh5*, which are presumed to act in the same pathway as MutLγ, results in failure to progress beyond early prophase I (Edelmann *et al.* 1999; de Vries *et al.* 1999; Kneitz *et al.* 2000; Santucci-Darmanin and Paquis-Flucklinger 2003). This suggests that MutSγ plays an essential role in early events of DSB repair and, indeed, studies in budding yeast have demonstrated a role for this complex in stabilizing DSB repair intermediates upstream of both NCO and CO resolution (Börner *et al.* 2004).

Confounding these observations are *in vitro* biochemical analyses which suggest that human MutSγ binds to dHJ substrates (Snowden *et al.* 2008), but these dHJs do not appear until pachynema of prophase I (Hunter and Kleckner 2001), well after the initial appearance of MutSγ. Unfortunately, it has not been possible to resolve the conflicting data in mouse spermatocytes *in vivo* because the loss of spermatocytes prior to zygonema in *Msh4^-/-^* or *Msh5^-/-^* males makes it impossible to investigate the importance of this complex for dHJ resolution and class I COs. More recent data indicate that the MutSγ of *S.* cerevisiae also binds HJs with strong affinity, with the additional ability to bind to early SEIs to stabilize these structures in advance of dHJ formation (Lahiri *et al.* 2018). Moreover, studies in mice show that MutLγ and MutSγ are present on SC during late pachynema, at a frequency and distribution that resemble class I CO (Santucci-Darmanin and Paquis-Flucklinger 2003). This suggests that, similar to other MMR complexes, MutSγ functions to recruit MutLγ to the SC during pachynema. At the same time, the earlier phenotype of MutSγ-deficient mice compared to MutLγ-deficient mice, coupled with the hyper-accumulation of MSH4 and MSH5 foci in zygonema (approximately 150 foci) compared to the limited recruitment of MLH1 and MLH3 in pachynema (approximately 25 foci), suggests additional functions of MutSγ from that of MutLγ.

All MutS proteins possess an ATPase domain that is essential for mismatch recognition, ATP binding, and for the initiation of mismatch excision during post-replicative mismatch repair (Bjornson and Modrich 2003; Mazur *et al.* 2006; Hingorani 2016; Liu *et al.* 2016). While MSH4 and MSH5 have lost the domain important for mismatch recognition, a similar ATP processing by human MSH4 and MSH5 has been proposed to play an important role in meiotic recombination in *in vitro* studies (Snowden *et al.* 2004, 2008), however, little is known about the significance of this for mammalian meiotic progression *in vivo*. To explore MSH5 ATPase function in greater detail we mutated a highly conserved residue within the P-loop domain of mouse *Msh5* (G to A mutation at residue 596, termed *Msh5*^GA^), which has been shown previously to affect ATP binding by MutS homologs. A similar mutation in *S. cerevisiae* revealed that loss of ATP binding in yeast Msh5 has no effect on the dimerization of mutant MSH5 with its wild-type (WT) MSH4 partner, but reduces crossing over and spore viability (Nishant *et al.* 2010). Based on this study and on the known yeast Msh4-Msh5 structure (Obmolova *et al.* 2000; Rakshambikai *et al.* 2013), we anticipated that the G-to-A mutation within the MSH5 ATP binding domain would not affect MutSγ complex formation. Interestingly, although spermatocytes in *Msh5^-/-^* mice fail to progress beyond zygonema, a subset of *Msh5^GA/GA^* spermatocytes escape this fate, progressing through prophase I and entering metaphase I. Thus, this mutant allele allowed for the first time, to investigate the role of MSH5 in crossing over during the entirety of mammalian prophase I. Interestingly, diakinesis-staged chromosomes from spermatocytes of *Msh5^GA/GA^* mice show exclusively univalent chromosomes and a complete absence of chiasmata, including those residual chiasmata that would presumably arise from the class II CO (MUS81-EME1) pathway. These observations indicate that the ATPase domain of MSH5 is essential for MutSγ activity early in DSB repair, and that mutation of this domain results in disrupted homolog interactions and aberrant DNA repair, leading to a failure to form any COs at the end of prophase I. Thus, loss of a functional MutSγ complex impacts CO formation regardless of the route of CO resolution.

## RESULTS AND DISCUSSION

### Generation of *Msh5^G596A^* mutant mice

In the current work, we generated a mouse line bearing a knock-in mutation that disrupts the conserved Walker “type A” motif GXXXX<u>G</u>KS/T (<u>G</u> refers to the modified G596 amino acid residue) in the ATPase domain of MSH5, which is important for ATP binding (Figure S1A). The targeting vector introduces a glycine to alanine change at amino acid residue 596 into exon 19 (Figure S1A,B). The mutant allele of this mouse line is designated *Msh5^G596^*^A^ (*Msh5^GA^*), and was predicted to abolish ATP binding in the MSH5 subunit. This mutation has previously been shown to preserve interaction with MSH4, allowing for the appropriate assembly of the MutSγ heterodimer. (Nishant *et al.* 2010). An additional diagnostic *BlpI* restriction site that does not alter the amino acid sequence was generated in the *Msh5* coding regions that overlaps with the mutation (Figure S1B). Transmission of the mutant *Msh5^GA^* allele was confirmed by PCR genotyping of genomic tail DNA and subsequent restriction of the associated *BlpI* site (Figure S1C,D). Likewise, RT-PCR and subsequent *BlpI* restriction digestion confirmed expression of the mutant transcript in *Msh5^+/GA^* and *Msh5^GA/GA^* mice (Figure S1D). In addition, while homozygous null mice lack all detectable MSH5 protein, the mutant MSH5^GA^ protein was expressed in the testis of both *Msh5^GA/+^* and *Msh5^GA/GA^* mice at levels similar to WT mice (Figure S1E). In all subsequent studies, *Msh5^GA/GA^* mutant animals were compared to *Msh5^+/+^* wildtype (WT) littermates, as well as *Msh4^-/-^* and/or *Msh5^-/-^* mice (Edelmann *et al.* 1999; Kneitz *et al.* 2000). All alleles of *Msh4* and *Msh5* were maintained on a C57Bl/6J background.

### *Msh5^GA/GA^* mice exhibit severely impaired meiotic progression, reduced testis size, and no spermatozoa

Similar to *Msh5^-/-^* and *Msh4^-/-^* mice, *Msh5^GA/GA^* mice are infertile as a result of defects in meiotic prophase I. By contrast, *Msh5^GA/+^* females and males are fertile (not shown), with no change in testis weights in *Msh5^GA/+^* males compared to WT littermates (Figure 1A). Homozygous mutant *Msh5^GA/GA^* males display a 40% reduced testis size compared to their WT littermates (Figure 1A) associated with complete loss of epididymal spermatozoa (Figure 1B). Sperm production in *Msh5^GA/+^* males was similar to that of WT animals (Figure 1B).

**Figure 1.**
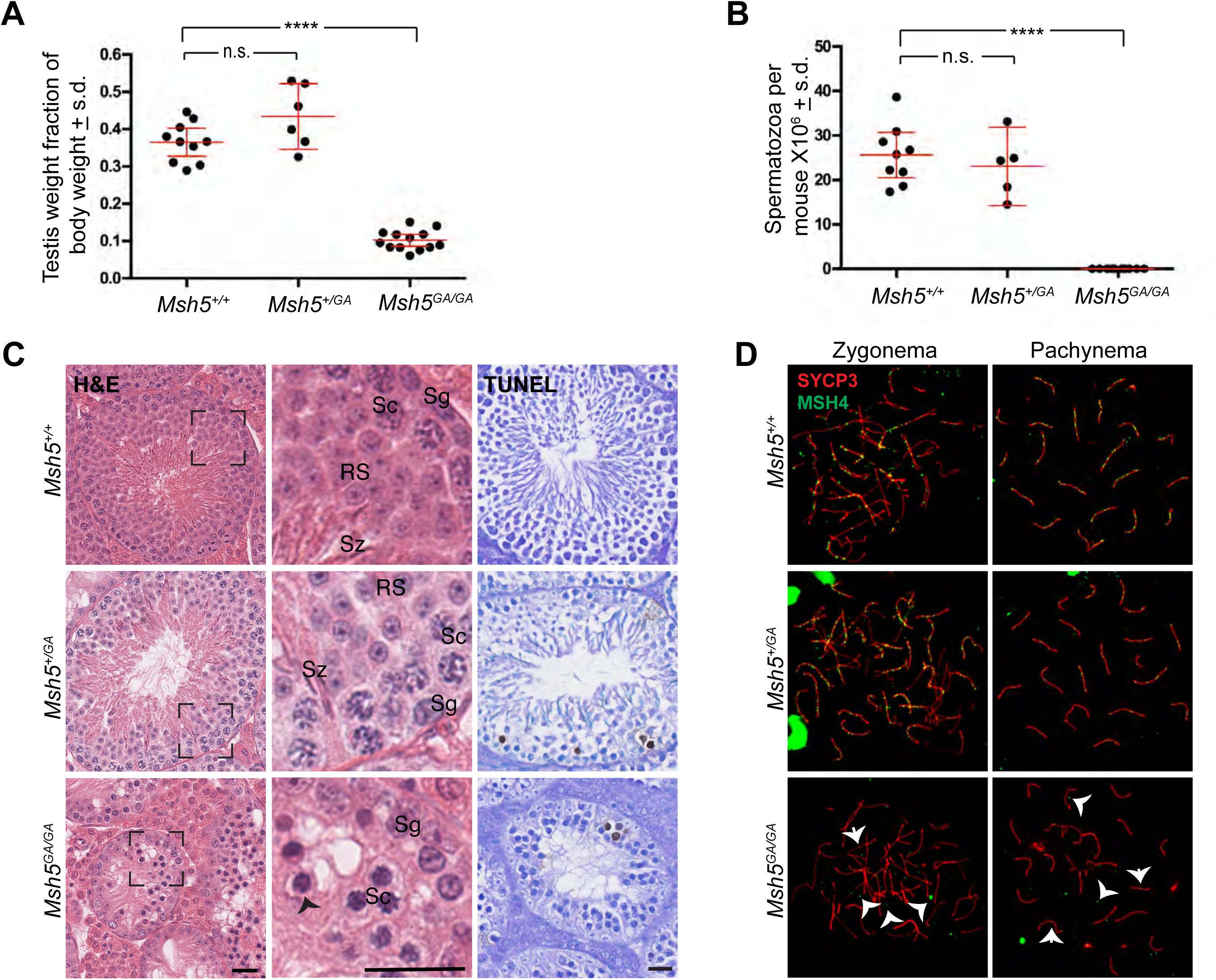
*Msh5^GA/GA^* mice confer an infertility phenotype that is not observed in Msh5^+/+^ or *Msh5^GA/+^* littermates. (a) Adult testis weights are significantly smaller in *Msh5^GA/GA^* mice compared to *Msh5^+/+^* littermates (p < 0.0001, unpaired t-test with Welch’s correction), while *Msh5^GA/+^* animals are not found significantly smaller than wild type littermates (*Msh5^GA/GA^* – 0.10% of total body weight ± 0.03, n = 13 testes; *Msh5^+/+^* −0.37% ± 0.05, n = 10; *Msh5^+/GA^* – 0.43% ± 0.08, n = 6; n.s.-not significant, **** p<0.0001). *Msh5^GA/GA^* animals have zero epididymal spermatozoa. (b) (WT – 17.4 × 10^6^ ± 6.6; p < 0.0001, unpaired t-test with Welch’s correction n = 9, n = 10; error bars show standard deviation). (c) Hematoxylin and Eosin staining of paraffin-embedded testis sections from *Msh5^+/+^*, *Msh5^+/GA^*, and *Msh5^GA/GA^* littermates. WT and heterozygous testes show meiotic and post-meiotic cells whereas *Msh5^GA/GA^* testes are absent of all spermatids and spermatozoa, and apoptotic cells are observed (Sg – spermatogonial, Sc – spermatocytes, RS – Round spermatids, Sz – Spermatozoa, arrow head shows apoptotic cell; scale bar represents 25nM). Boxes represent magnified image on the right. TUNEL assay reveal apoptotic cells within seminiferous tubules. (d) MSH4 (green) staining on chromosome spreads for adult *Msh5^+/+^* and *Msh5^+/GA^* zygotene and pachytene spermatocytes localize MSH4 to the SC (red) during zygotene and pachytene, while the association between MSH4 and the SC is largely disrupted in *Msh5^GA/GA^* spermatocytes. Arrow heads in zygotene panel highlight common localization of MSH4, with few foci associated with the SC and frequent foci of varying sizes localized away from the SC. Arrow heads in mutant pachytene panel point out faint MSH4 signal on or near the SC suggesting the association between MSH4 and SC is not entirely abolished in the mutants.

Hematoxylin and Eosin (H&E) staining of testis sections from WT adult male mice showed normal cell populations within the seminiferous epithelium, while spermatogenesis was severely disrupted in *Msh5^GA/GA^* testis sections (Figure 1C). Testis sections from *Msh5^GA/GA^* male mice contained Leydig cells, Sertoli cells, and spermatogonia, and spermatocytes, along with a high proportion of TUNEL-positive apoptotic germ cells within the seminiferous tubules (Figure 1C). Most notably, testis sections from *Msh5^GA/GA^* males contained numerous pachytene and post-pachytene spermatocytes (Figure 1C, arrow), including cells that were clearly at metaphase I. This is in contrast to our previous observations in *Msh5^-/-^* and *Msh4^-/-^* males, in which the entire spermatocyte pool is lost at or prior to entry into pachynema ((Edelmann *et al.* 1999; Kneitz *et al.* 2000) and Figure S2). Seminiferous tubules from WT males have an average of less than one TUNEL-positive cell while the heterozygous *Msh5^GA/+^* tubules have an average of 1 TUNEL-positive cell per tubule. The *Msh5^GA/GA^* tubules have a higher average of 2.7 TUNEL-positive cells per tubule. The important difference between the histological appearance of *Msh5^GA/GA^* tubules and that of *Msh5^-/-^* and *Msh4^-/-^* males is the increased progression into pachynema and the appearance of numerous metaphase cells in the tubules of *Msh5^GA/GA^* males.

### A functional ATP binding domain of MSH5 is important for early homolog interactions and complete homolog synapsis

To assess progression through prophase I, immunofluorescence (IF) staining was performed on chromosome spreads of spermatocytes from *Msh5^+/+^* and *Msh5^GA/GA^* adult male mice using antibodies against components of the SC, SYCP3 and SYCP1 (Figure 2A). In leptonema, the SC begins to form, with SYCP3 localization appearing in a punctate pattern along asynapsed chromosomes. Such a staining pattern was evident on leptotene chromosome preparations from mice of all genotypes (Figure 2A). Upon entry into zygonema, the transverse filaments and central element of the SC begins to assemble, as shown by the localization of SYCP1 between the chromosome axes on chromosome spreads from both *Msh5^+/+^* and *Msh5^GA/GA^* adult male mice (Fig 2A). By pachynema, when autosomes in *Msh5^+/+^* adult males are now fully synapsed along their entire lengths, the first signs of synapsis failure become evident in *Msh5^GA/GA^* animals. While WT pachytene cells contain 20 independently synapsed homologs, spermatocytes from *Msh5^GA/GA^* animals show variable degrees of synapsis, coupled with frequent occurrences of inappropriate synapsis between non-homologous partners (Figure 2A, arrowheads), synapsis between multiple chromosomes, but also some occurrences of apparently normal homolog synapsis. The aberrant synapsis phenotype observed in *Msh5^GA/GA^* spermatocytes ranges in severity, with some pachytene-like cells showing synapsis defects across the majority of homolog pairs, while other pachytene-like cells showed defects among a few homolog pairs.

**Figure 2.**
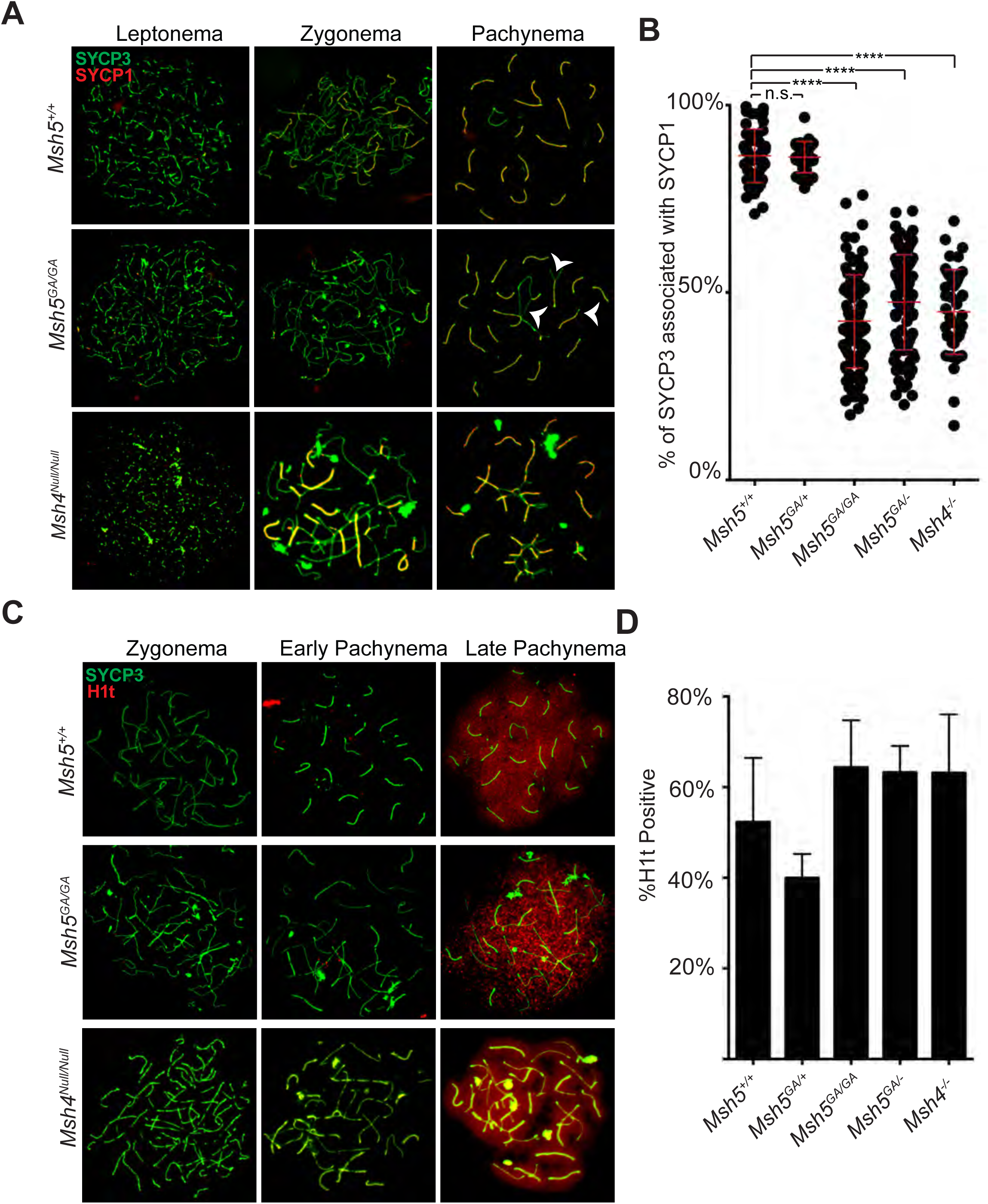
*Msh5^GA/GA^* spermatocytes have inappropriate synapsis between non-homologous chromosomes and progress through mid-pachytene. Localization of lateral element SYCP3 (green) (a,c) with the localization of central element protein SYCP1 (red) (a) in adult chromosome spreads show the normal progression of synaptonemal complex through prophase I in *Msh5^+/+^* spermatocytes. *Msh5^GA/GA^* spermatocytes have varying degrees of synapsis during mid-prophase I with notable inappropriate synapsis between multi-homologs associations (arrow heads). (b) Synapsis was measured by comparing lengths of total SYCP3 tracks to the total lengths of SYPC1 tracts in each pachytene-like cell using Image J. Percentage of synapsis is calculated per cell by dividing SYCP1 length with SYCP3 length and multiplying by 100. Each point represents a different pachytene-like cell, red overlay lines depict the average ± SD. On average, *Msh5^+/+^* pachytene spermatocytes had 86% synapsis. Compared to wild-type, *Msh5^GA/+^* animals are not found to be significantly different and had an average of 78% synapsis (unpaired t-test with Welch’s correction, p<0.47). *Msh5^GA/GA^, Msh5^GA/-^, and Msh4^-/-^* animals are all found to be significantly different from wild-type (unpaired t-test with Welch’s correction, p < 0.0001, p < 0.0001, and p < 0.0001 respectively). Average synapsis for *Msh5^GA/GA^, Msh5^GA/-^, and Msh4^-/-^* pachytene-like cells was 42%, 47%, and 45% respectively. (c) Localization of mid pachytene histone marker, H1t (red) is observed on both *Msh5^+/+^* and *Msh5^GA/GA^* mid pachytene to diplotene spermatocytes. (d) % H1t positive prophase I stages are shown for each genotype (late pachytene and diplotene populations). When compared to wild-type, the *Msh5^GA/GA^, Msh5^GA/-^, and Msh4^-/-^* animals all had comparable H1t positive populations across prophase I (unpaired t-test with Welch’s correction, p < 0.1143, p < 0.1714, and p <0.3243 respectively).

Utilizing Image J software, we were able to obtain quantitative measurements of synapsis across our mouse model. For each cell we measured the total track length of SYCP3 signal and compared it to the total track length of SYPC1 to obtain the percent synapsis (SYCP1/SYCP3 × 100). For this analysis, we used *Msh4^-/-^* mice as a comparison with *Msh5^GA/GA^* males because the original reports suggested slightly higher levels of synapsis than observed in *Msh5^-/-^* mice, and because *Msh5^-/-^* mice are no longer available. Since MSH4 and MSH5 always act as a heterodimer, these mouse mutants reflect overall MutSγ function. Previous descriptions of *Msh4^-/-^* males indicated no pachytene entry, an observation that was based on the 20 independently synapsed homologs as defined by WT pachytene. In the current study however, we defined “pachytene-like” as >4 synapsed or partially synapsed chromosomes. Under these criteria, we observe pachytene-like cells in both *Msh4^-/-^* males and in *Msh5^GA/GA^* males.

The average synapsis in WT spermatocytes during pachynema, remembering the XY chromosome pair in males is only synapsed at the autosomal region, is 86.5 ±

7.2% (Figure 2B) with *Msh5^GA/+^* spermatocytes showing normal synapsis 86.3 ± 4.2% (Figure 2B). Overall there is a remarkable degree of synapsis in *Msh5^GA/GA^* animals, with spermatocytes exhibiting an average of 43.2 ± 12.4%, and some cells achieving up to 76% of chromosome axes. By contrast, synapsis in *Msh5^-/-^* animals is less than 5% in two previous reports ((Edelmann *et al.* 1999) and Figure 2B). The level of synapsis in *Msh5^GA^*^/-^ males is comparable to that of *Msh5^GA/GA^* males, at 47.4 ± 12.7% synapsis. Synapsis in *Msh4^-/-^* males was slightly lower than *Msh5^GA/GA^* males, at 44.9 ± 11.3% synapsis (Figure 2B). Importantly, synapsis in *Msh5^GA/-^* spermatocytes is similar to that seen in *Msh5^GA/GA^* homozygous mutant animals, while synapsis in *Msh5^GA/+^* spermatocytes is similar to WT, indicating that the *Msh5^GA^* allele is recessive and not causing a dominant negative effect.

The histone marker, H1t, allows for differentiation of pachytene cells into “early” and “late”, since H1t only associates with the latter population (Wiltshire *et al.* 1995). Despite the incomplete synapsis and inappropriate synapsis between multiple chromosomes in spermatocytes from *Msh5^GA/GA^* animals, these cells are competent to achieve a mid-pachytene-like stage of meiosis, at least as assessed by acquisition of H1t signal (Figure 2C). Synapsis mutants (*Msh5^GA/GA^*, *Msh5^GA/-^*, *Msh4^-/-^*) do not achieve the normal 20 independently synapsed homologs as observed in WT. However, the localization of H1t to these mutants suggests that they are achieve a pachytene-like stage. Alternatively, this could suggest that the meiotic progression is disjointed between cell processes.

To compare prophase I populations across genotypes, we looked at the total number of H1t-positive cells in prophase I (Figure 2D). Across all prophase I cells in WT males, we observe that 52.3 ± 14.1% of cells are H1t-positive. Surprisingly, our mutant animals gave values similar to WT: in *Msh5^GA/GA^* animals we observe a 64.4 ± 10.1% H1t positive prophase I population, in *Msh4^-/-^* 63.2 ± 12.9%; *Msh5^-/GA^* 63.3 ± 5.8%. The *Msh5^+/GA^* spermatocytes are the only population we observed a lower, albeit not statistically different H1t-positive prophase I pool of 40.0 ± 5.3%. Overall, we observe a comparable prophase I progression in *Msh5^GA/GA^* mutant spermatocytes and in *Msh4^-/-^* spermatocytes.

The results presented here are in agreement with first reports of *Msh4* which showed cell death to occur at prophase I. Our results were able to further narrow down the window of cell death for *Msh4* spermatocytes to pachytene-late diplotene. This is reasonable given we employed tools that were not available at the time of the original report on the *Msh4^-/-^* allele. Specifically, synapsis quantitation using Image J, and the histone H1t marker

To further assess the degree of synapsis in different mice, the number of independently synapsed homologs were counted in each pachytene-like cell across the *Msh4^-/-^* and *Msh4^-/-^* males (Figure S3). Neither the *Msh4^-/-^* males nor the *Msh4^-/-^* males are able to achieve a wild-type pachytene configuration of 20 independently synapsed homologs. While some *Msh4^-/-^* pachytene cells were only able to achieve as many as 12 independently synapsed homologs, only 4.8% of the pachytene-like population had more than 10 independently synapsed homologs. The degree of synapsis is greater in *Msh5^GA/GA^* and *Msh5^GA/-^* spermatocytes, with instances of cells achieving up to 15 independently synapsed homologs occurring in each genotype, and *Msh5^GA/GA^* having 5.6% homologs having more than 10 independently synapsed homologs and *Msh5^GA/-^* having 11.6% homologs having more than 10 independently synapsed homologs (Figure S3).

The greater degree of synapsis observed in spermatocytes from *Msh5^GA/GA^* or *Msh5^GA/-^* males compared to that of *Msh4^-/-^* cells suggests that the presence of the MSH5^GA^ protein allows for more proficient early homolog pairing, or that SC is established more robustly in the presence of defective MutSγ heterodimer than in the complete absence of any heterodimer. Thus, homolog pairing and/or synapsis initiation/progression does not rely on a fully functional MutSγ heterodimer.

### MutSγ association with the synaptonemal complex is drastically reduced in spermatocytes from *Msh5^GA/GA^* males

In WT mice, MSH4 and MSH5 localize on chromosome cores of the SC from zygonema through pachynema, with approximately 200 foci in zygonema, reducing progressively through until late pachynema (Kneitz *et al.* 2000). We investigated whether MutSγ localization on SCs was affected by loss of a functional ATP binding domain within MSH5. To this end, chromosome spreads from *Msh5^+/+^*, *Msh5^+/GA^*, and *Msh5^GA/GA^* male mice were subjected to IF staining using antibodies against MSH4 and SYCP3. MSH4 localization in early prophase I cells look comparable between *Msh5^+/+^* and *Msh5^+/GA^* adult males, with abundant foci associated with early SC structures in zygonema (Figure 1D). Interestingly, in spermatocytes from *Msh5^GA/GA^* males, there appears to be a dramatically decreased association of MSH4 to the SC and an observable increase in MSH4 foci not associated with the SC in zygotene and pachytene nuclei (Figure 1D). Overall the intensity of MSH4 staining in zygotene and pachytene spermatocytes from *Msh5^GA/GA^* males is lower than that of WT littermates, although some foci are clearly associated with the SC at both zygonema and pachynema (Figure 1D, arrows). Further examples of MSH4 staining at this stage are provided in Figure S4 which provide additional evidence of a broader but fainter distribution of MSH4 signal in spermatocytes from *Msh5^GA/GA^* males. MSH4 localization away from the SC may indicate that the MSH5 ATP binding domain is essential for the recruitment and retention of MutSγ with the SC cores from zygonema through until pachynema. Alternatively, this reduction of MSH4 signal could be due to destabilization of the mutant MutSγ heterodimer followed by degradation of MSH4 protein. Importantly, though some MSH4 is associated with chromosome cores when bound to the MSH5^GA^ protein, this localization is greatly diminished compared to MutSγ localization in WT spermatocytes.

### The ATP binding domain of MSH5 is essential for timely progression of DSB repair events

To assess progression of DSB repair through prophase I, IF was performed on chromosome spreads from *Msh5^+/+^*, *Msh5^GA/GA^*, and *Msh5^+/GA^* adult littermates using antibodies against γH2AX, a phosphorylated histone variant that marks sites of DSB (Figure 3A). Spermatocytes from *Msh5^+/+^* animals show a strong γH2AX signal during leptonema and zygonema of prophase I indicating normal induction of DSBs, with loss of the γH2AX signal at pachynema signaling progression of DSB repair (Figure 3A). As expected, the γH2AX signal is intensified on the sex chromosomes at pachynema, a phenomenon that is not related to DSB formation (Turner *et al.* 2004); (Figure 3A, top row). Spermatocytes from *Msh5^GA/GA^* animals show a similarly strong γH2AX signal during leptonema and zygonema, indicating DSBs are induced at the expected time. Unlike in *Msh5^+/+^* cells, however, γH2AX signal is retained on autosomes throughout prophase I in *Msh5^GA/GA^* cells, indicating persistent DNA damage (Figure 3A, bottom row).

**Figure 3.**
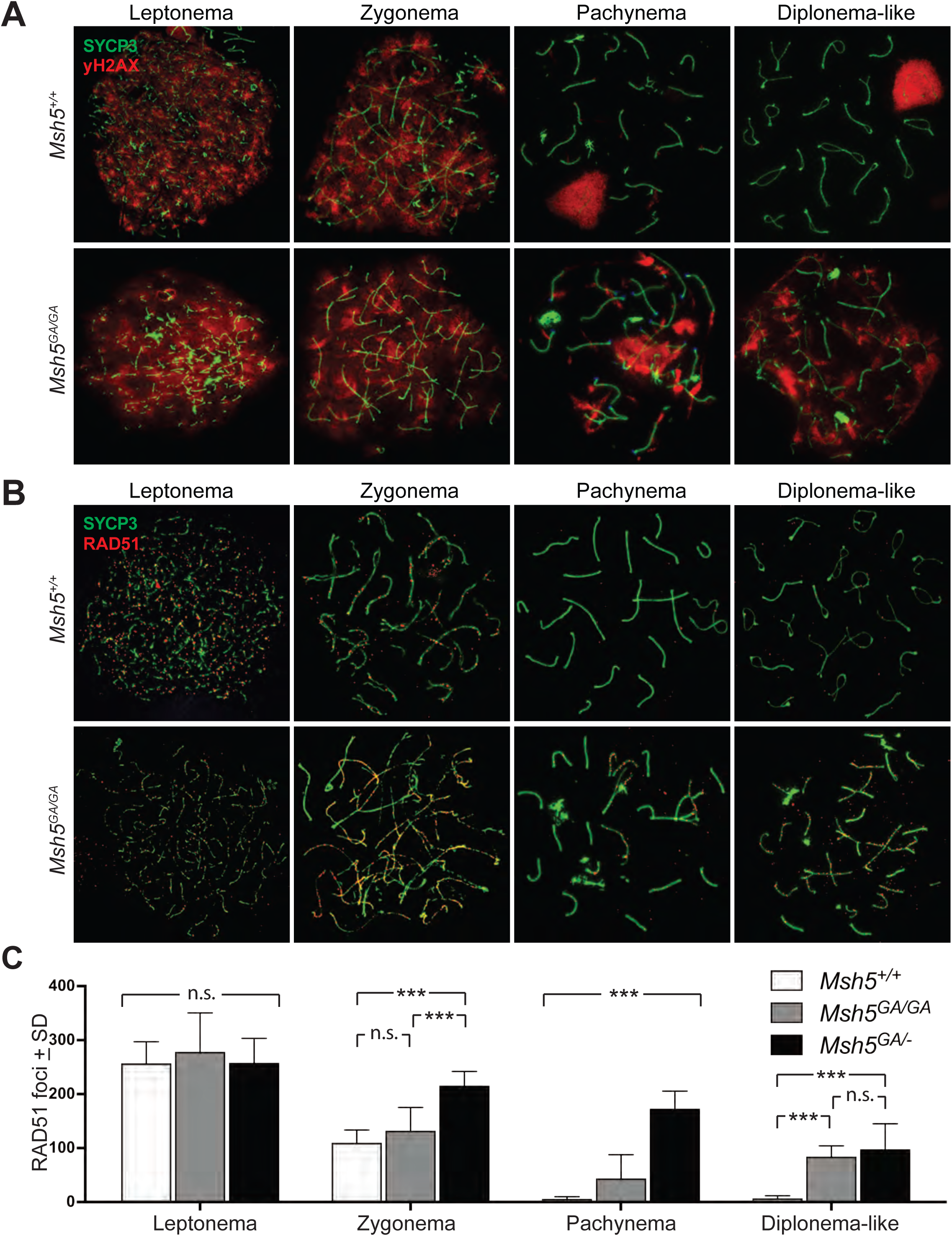
DNA damage persists in *Msh5^GA/GA^* spermatocytes throughout prophase I. (a) Immunofluorescent staining of yH2aX (red) on chromosome spreads of *Msh5^+/+^* and *Msh5^GA/GA^* littermates. (b) DNA repair marker RAD51 (red) on *Msh5^+/+^* and *Msh5^GA/GA^* chromosome spreads persists throughout prophase I. (c) Quantification of RAD51 foci associated with the SC of chromosome spreads during leptonema (*Msh5^+/+^*-257.2 ± 39.74, n = 13; *Msh5^GA/GA^* −278.6 ± 71.92, n = 14; p=0.88, Mann-Whitney) zygonema (*Msh5^+/+^* −109.9 ±23.72, n=30; *Msh5^GA/GA^* −132.4±43.04, n=29; p=0.14, Mann-Whitney), pachynema (*Msh5^+/+^* −6.11 ± 4.12, n=26; *Msh5^GA/GA^* −97.66 ± 43.84, n = 35; p<0.0001, Mann-Whitney) and diplonema (*Msh5^+/+^* −6.91 ± 4.85, n=11; *Msh5^GA/GA^* −88.18 ± 20.02, n=11; p <0.0001, Mann-Whitney)

During DSB repair, one of the earliest common intermediate steps involves strand invasion and homology search, which is mediated by the RecA homologs, RAD51 and DMC1. MutSγ has been suggested to participate in stabilization of these strand invasion events *in vitro* (Snowden *et al.* 2004). During leptonema in WT spermatocytes, RAD51 foci are observed on axial elements of the SC in high numbers (Figure 3B,C), and similar numbers of RAD51 foci are observed on leptotene spreads from *Msh5^GA/GA^* spermatocytes. As WT cells progress from Zygonema to pachynema, RAD51 foci numbers drop dramatically, reflecting the repair of DSBs.

The RAD51 focus numbers in *Msh5^GA/GA^* and *Msh5^GA/-^*spermatocytes remain significantly elevated above that of WT spermatocytes throughout prophase I (p<0.0001, Figure 3C). Interestingly, the RAD51 focus counts at zygonema and pachynema are significantly higher in *Msh5^GA/-^* spermatocytes than in homozygous mutant *Msh5^GA/GA^* spermatocytes (p<0.0001, Figure 3C), indicating more DSB repair activity during this stage in the presence of only one copy of ATPase defective *Msh5.* At diplonema, WT spermatocytes have lost all RAD51 foci, but these foci remain significantly higher in *Msh5^GA/GA^* and *Msh5^GA/-^* spermatocytes, albeit at lower frequency to that seen in pachynema (p<0.0001). At this stage, RAD51 counts in *Msh5^GA/GA^* and *Msh5^GA/-^* spermatocytes are not statistically different from each other. Importantly, spermatocytes from *Msh5^GA/+^* males are similar to WT with few abnormalities and normal dynamics of RAD51 loss (Figure 3C, S3B).

Taken together, these data demonstrate that normal DSB induction occurs in the presence of the ATP binding-defective MSH5^GA^ protein, but DSB repair is disrupted. Alternatively, it is possible that the high rate of RAD51 foci observed at pachynema in *Msh5^GA/GA^* males results from additional induction of DSBs through prophase I, but the current tools preclude our ability to differentiate between these two options. Importantly, the presence of only one GA allele on a WT background (*Msh5^GA/+^* males) results in normal temporal dynamics of RAD51 loss, while the presence of one GA allele on a null background (*Msh5^GA/-^* males) results in a significantly more delayed processing of DSBs, as characterized by RAD51 accumulation and loss. These observations argue strongly against a dominant negative effect of the GA point mutation.

### An intact MSH5 ATP binding domain is essential for formation of all classes of crossover

MutSγ recruits the MutLγ complex during pachynema as part of a canonical class I CO machinery. IF staining using antibodies against MLH3 was compared across genotypes (Figure 4A). In WT and *Msh5^GA/+^* mice during pachynema, MLH3 appears on chromosome cores at a frequency that correlates with final class I CO numbers (Figure 4A, top row), but is absent at in spermatocytes from *Mlh3^-/-^* males (Figure S5B). In pachytene-like spermatocytes from *Msh5^GA/GA^* males, MLH3 foci do not form on chromosome cores (Figure 4A). Occasional very faint signal was observed throughout the chromatin, as well on the SC cores, when the microscope intensity gain is increased, but it was not possible to obtain reliable images depicting this weak signal. Nonetheless, such staining was never observed in chromosome spread preparations from *Mlh3^-/-^* males, suggesting that this weak staining might be specific for MLH3 protein (Figure S5B). Given the diffuse and faint nature of this staining, it cannot be determined if this MLH3 signal is associated with sites of DSB repair. Thus, a fully functional MSH5 protein is required for appropriate association of the MutLγ with the synaptonemal complex and establishment of nascent class I CO sites.

**Figure 4.**
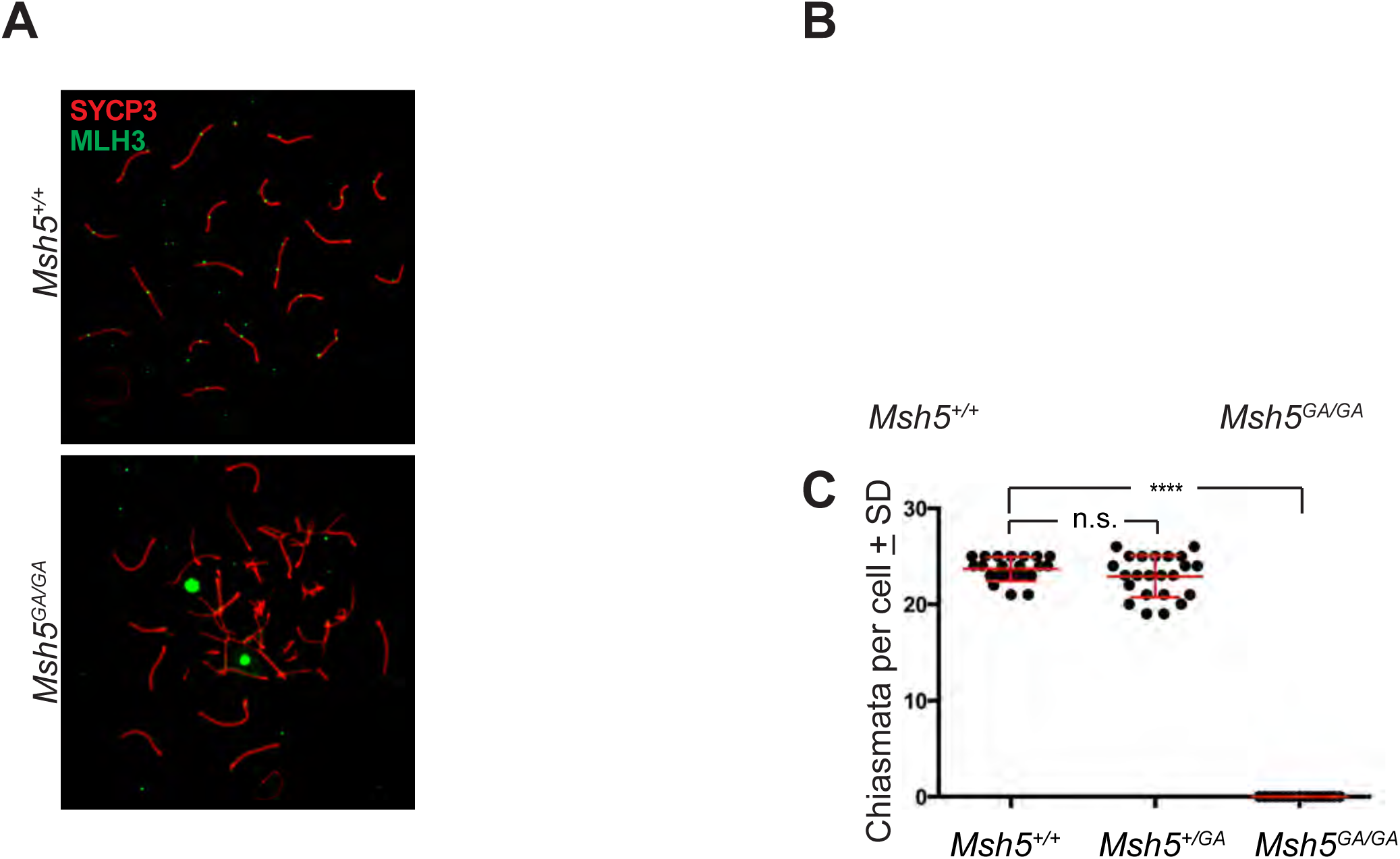
No crossovers form in *Msh5^GA/GA^* spermatocytes. (a) Immunofluorescent staining of MLH3 (green) in adult pachytene *Msh5^+/+^* and *Msh5^GA/GA^* chromosome spreads show localization of MLH3 to SC (red) as expected in wild type and no MLH3 localization to the SC in mutant. (b) Giemsa staining of diakinesis preparations from *Msh5^+/+^* and *Msh5^GA/GA^* litter mates showing normal chiasmata in wild type and completely univalent chromosomes in mutants. (c) Chiasmata counts for *Msh5^+/+^* (23.68 ± 1.25, n=22) and *Msh5^GA/GA^* (0.0, n = 15) litter mates (p<0.0001, unpaired t-test).

To assess crossing over across the genome, diakinesis spreads were prepared to assess chiasmata formation (Holloway *et al.* 2010). In WT males, each bivalent chromosome pair has at least one chiasmata (Figure 4B). Since a small number of spermatocytes from *Msh5^GA/GA^* males are capable of progressing into diakinesis, we were able to count chiasmata in these homozygous mutant mice (Figure 4B,C). Unexpectedly, diakinesis-staged cells from *Msh5^GA/GA^* males displayed exclusively univalent chromosomes and did not form any chiasmata (Figure 4B,C). Thus, normal MSH5 ATP processing is essential for all crossover formation in mammals. Such analysis has not been possible in *Msh5^-/-^* males because spermatocytes for these mice fail to reach diakinesis. Taken together, these results demonstrate that MutSγ function is essential for all CO events, regardless of the pathway by which they were created.

### ATP processing activity in MSH4 alone is insufficient to promote full MutSγ activity

Each of the ATPase domains of a canonical MutS complex (MutSα, MutSβ) are essential for mismatch recognition through ADP/ATP binding and hydrolysis (Haber and Walker 1991; Gradia *et al.* 1997; Acharya *et al.* 2003; Kijas *et al.* 2003). In MutSγ complex, the MSH5 has been shown to bind ATP with a higher affinity than MSH4 suggesting asymmetric ATP activity among MutSγ subunits (Snowden *et al.* 2008). The work presented here suggests that the ATPase domains from both MSH4 and MSH5 subunit proteins are also utilized by MutSγ *in vivo*. Our analysis of the *Msh5^GA/GA^* allele has shown that in the MSH5^GA^-MSH4 complex, the MSH4 ATP binding domain by itself is insufficient for the complete repertoire of MutSγ functions including both the early functions essential for early homolog interactions, as well as later functions in CO processing.

*In vitro* biochemical studies have demonstrated that, in the absence of ATP, the human MutSγ complex encircles Holliday junctions forming a clamp around the junction of two DNA helices. Upon ATP binding, the MutSγ clamp slides along the two DNA helices, moving away from the initial Holliday junction (Snowden *et al.* 2004). From this work it was proposed that the exchange of ADP to ATP results in the formation of an ATP hydrolysis-independent MutSγ sliding clamp, and that as each MutSγ moves away from the Holliday junction core, it allows subsequent loading of additional MutSγ complexes which encircle two homologous duplex DNA arms, thus stabilizing the paired DNA double strands (Bocker *et al.* 1999; Snowden *et al.* 2004, 2008). While it should be noted, the *in vivo* substrate has yet to be determined for MutSγ, but given the timing of MutSγ foci appearance around the time of RAD51, SEI is a candidate for an initial substrate. Indeed, recent *S. cerevisiae* data support this conclusion (Lahiri *et al.* 2018).

Loss of ATP binding in our mutants is predicted to result in a clamp protein that is unable to slide. In our *Msh5^GA/GA^* mutants we see a large reduction in MSH4 signal along chromosome cores, but some MSH4 persists, suggesting either minimal loading of MutSγ complex onto the DNA and/or enhanced (but not complete) degradation of the complex. One possibility is that the MSH5^GA^-MSH4 clamp protein is still able to provide some stabilization between homologs, allowing for small amounts of synapsis in *Msh5^GA/GA^* animals. However, without the sliding ability, the normal function of MutSγ in SC establishment and/or DSB repair processing is abolished.

### MutSγ loads onto a large number of DSB repair intermediates and is essential for all COs regardless of the pathway by which they are produced

In yeast, MutSγ has been shown to work in coordination with MutLγ to aid in the repair of DSBs at crossover events through the class I (ZMM) pathway. In the mouse, MutSγ accumulation on SCs in zygonema is in excess of the final number of MutLγ foci, but the two heterodimeric complexes are shown to localize similarly by late pachynema (Novak *et al.* 2001; Santucci-Darmanin and Paquis-Flucklinger 2003). In WT mouse spermatocytes, the earlier and more abundant localization of MutSγ in zygonema implies that MutLγ is recruited to only a subset of MutSγ sites upon entry into pachynema, with the remaining sites that fail to accumulate MutLγ presumably being processed to become NCO events via other repair pathways. Thus, the higher numbers of MutSγ foci in zygotene and early pachytene mouse spermatocytes, together with the earlier loss of spermatocytes in *Msh5^-/-^* animals compared to *Mlh3^-/-^* or *Mlh1^-/-^* mice, implies a role for MSH4 and MSH5 in DSB processing at an early intermediate stage for multiple repair pathways. Such a possibility is supported by our data showing that diakinesis preps from *Msh5^GA/GA^* spermatocytes display no chiasmata (Figure 4), which indicates that a functional MutSγ complex is essential for all CO, acting at a stage that is upstream of both class I and class II CO designation, and thus may be a common intermediate for all CO pathways early in prophase I. Indeed, both Class I and Class II crossovers arise from a common DNA repair intermediate structure downstream of RAD51/DMC1 activity. In *Msh5^GA/GA^* spermatocytes, the retention of RAD51 on SC late into pachynema suggests that DNA repair is delayed prior to either crossover pathway, though other possible explanations exist. Thus, while the class II CO pathway, which involves at least one structure specific endonuclease, MUS81-EME1 (Holloway *et al.* 2008; Schwartz and Heyer 2011), is not traditionally viewed to be dependent on the ZMM class of proteins, our data indicate that a functional MSH5 protein is required to promote both classes of CO. Conversely, while we briefly considered the possibility that the mutant MutSγ complex may bind irreversibly to DSB repair intermediates that might otherwise have been processed via the Class II pathway, thus blocking the recruitment of appropriate class II repair factors, this does not appear to be the case since little to no MSH4 staining is observed on the SC. The diffuse staining of MSH4 away from chromosome cores suggests that loss of appropriate loading of MutSγ on the SC is sufficient to prevent any CO processing, regardless of the pathway of repair.

The results provided here argue against the current dogma that states that ZMM proteins, of which MSH4 and MSH5 are family members, do not operate outside of the class I machinery. However, while our current data do not currently provide a mechanism by which MutSγ can orchestrate both CO pathways in mammals, studies from other organisms provide interesting insight into potential mechanisms. In *Tetrahymena thermophila*, for example, which has no SC, COs are exclusively of the class II variety, requiring Mus81-Mms4, but not the canonical ZMM family. Despite the absence of class I CO events, MSH4 and MSH5 are essential for appropriate CO levels in this species, leading to the conclusion that these proteins function outside (or upstream) of the canonical class I CO pathway (Shodhan *et al.* 2014).

In SC-bearing organisms, where class I and class II CO events occur in tandem to differing degrees, ZMM proteins appear to function exclusively in the metabolism of the former class of COs. In *S. cerevisiae*, CO assignment occurs prior to SC assembly, and the number of MSH5 foci observed in this species corresponds well with the final tally of class I COs (Agarwal and Roeder 2000). However, this does not appear to be the case for organisms such as *C. elegans,* in which only class I COs occur. Yokoo *et al* have proposed the installation of MSH-5 in worms represents a “CO licensing” stage during which the protein initially accumulates at a supernumerary frequency along the chromosome cores (Yokoo *et al.* 2012). These foci then diminish in number as the cell progresses through pachynema in *C. elegans*, accumulating the pro-crossover factor COSA-1 only once the final number of class I events is achieved. Thus, the final appearance of COSA-1 and MSH-5 bound foci at six sites across the worm genome represents the final “designation” of presumptive class I CO sites (Yokoo *et al.* 2012). Importantly, the accumulation of these pro-CO factors at DSB sites is postulated to result in a change in SC status, resulting in enrichment/stabilization of SC proteins at these sites, presumably favoring crossing over at these locations (Pattabiraman *et al.*). This would suggest that alterations in localized SC architecture in response to deposition of pro-CO factors may promote DSB repair towards a CO fate in worms. These distinct differences in CO designation and MutSγ installation are depicted in Figure 5.

**Figure 5.**
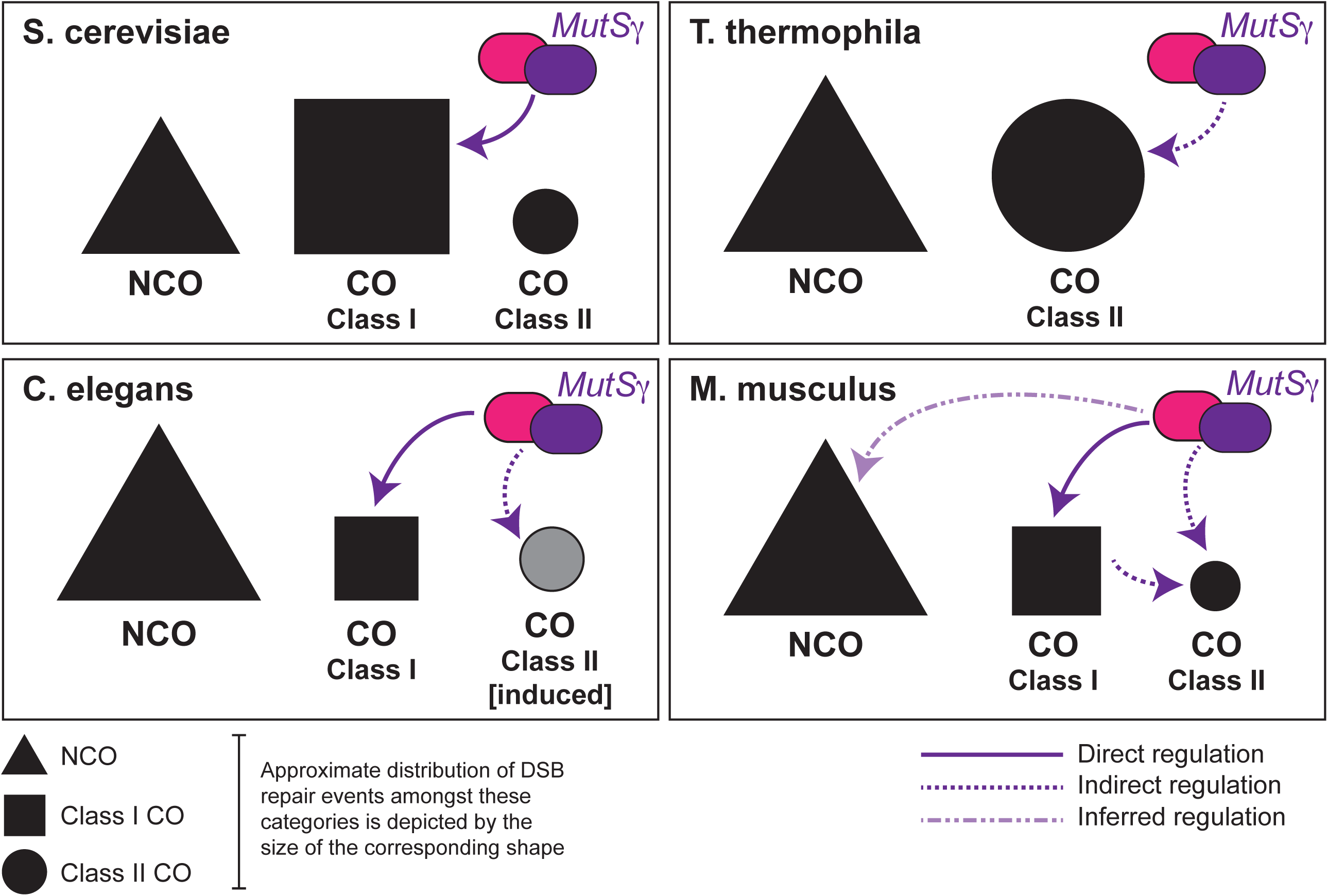
Model for MutSy function in different meiotic species.

In the mouse, the same excessive number of MutSγ foci appear somewhat earlier in prophase I, at or soon after the completion of the axial elements in early zygonema, and these too get pared down through zygonema and pachynema coincident with the progression of CO designation. Loss of the entire MSH5 protein results in a failure to accumulate MutSγ or to complete synapsis in zygonema, resulting in cell death prior to pachynema or, at the very most, aberrant progression through pachynema (Edelmann *et al.* 1999; Kneitz *et al.* 2000). These studies imply that, in the mouse at least, CO licensing is tightly linked to appropriate synapsis and may reflect the requirement for distinct rearrangements in SC architecture by the MutSγ complex, similar to that proposed for *C. elegans* (Pattabiraman *et al.*). However, in the current studies, we find that loss of a functional ATPase domain in one component of MutSγ, MSH5, allows for partial synapsis implying that any structural changes to the SC can be orchestrated in the absence of full ATPase activity of the MutSγ complex. Under such circumstances we lose all COs, regardless of their final pathway of biogenesis. Thus, CO processing through both the class I and class II pathways is dependent on a fully functional MutSγ heterodimer, but may not be dependent on any SC changes induced by MutSγ in zygonema.

Our data suggest either that functional activity of class II machinery depends on presence and processing of class I COs (an indirect requirement perhaps involving more discrete localized changes in the SC state at the DSB site), or that loading of class II pathway mediators requires the presence of MutSγ at these sites (a direct requirement for loading of MutSγ prior to recruitment of class II repair factors). In either case, this would infer that MutSγ is required for CO licensing for both pathways and/or lies upstream of the licensing decision. This is not surprising given that, in the mouse, no fewer than 60% of the DSB sites become loaded with MutSγ (or 150 out of 250), and only a minor fraction of these licensed sites (approximately 20%) will become COs of the class I or class II variety (Kneitz *et al.* 2000; Cole *et al.* 2012). Thus, there is an over-abundance of available sites for crossing over and, indeed MutSγ loads as efficiently onto NCO-destined DSB repair intermediates as it does onto CO-destined DSB repair intermediates. Though the implications of this promiscuous MutSγ binding is not yet understood, it suggests that, while CO licensing in worms is achieved by MSH-5 association, this may not be the case in the mouse since MutSγ association with DSB repair intermediates appears to be more promiscuous than in worm and yeast.

Taken together, our analysis of a point mutant mouse for *Msh5* has allowed us for the first time to explore late prophase I roles for MSH5 in DSB repair and homologous recombination. Our observations demonstrate that the large number of MutSγ sites found in early prophase I may serve as intermediates for both class I and class II CO events, and indeed for NCO events. In addition, we propose that MutSγ may function as a point of dialog between the two classes of CO event: notably, the loss of class I events does not disrupt class II CO (Edelmann *et al.* 1996; Lipkin *et al.* 2002; Svetlanov *et al.* 2008), whereas loss of class II CO results in a compensatory increase in class I CO, with additional CO events no longer being constrained by interference (Holloway *et al.* 2008). Thus, our observation that loss of functional MutSγ results in loss of both class I and class II COs at the end of prophase I could be indirect through direct actions on class I COs or, more likely, is due to the pre-loading of MutSγ at all DSB repair intermediates. Along these lines, it is important to note that the total number of MutSγ foci is around 150 in mid-zygonema, a number that is almost 6-fold higher than the total CO frequency in male mice. It is likely, therefore, that the majority of DSB repair events that become loaded with MutSγ are processed via NCO pathways including, but not limited to, SDSA. Moreover, unlike the situation in yeast, where NCO events are not associated with MutSγ loading and are all processed within a similar time frame, the loading of MutSγ in mouse spermatocytes results in progressive NCO formation through prophase I. Thus, MutSγ loading in mouse meiosis does not restrict DSB repair to a class I CO fate. Instead, loading of mammalian MutSγ can lead to NCO or CO fates, the latter involving multiple classes of CO. Given that MutLγ is restricted to class I CO events, these data suggest a functional distinction between the roles of MutSγ and MutLγ in DSB repair during mammalian meiosis, and open the door for additional roles for MutSγ in orchestrating/overseeing DSB repair in the mammalian germline. In light of the role of other heterodimeric MutS complexes in recruiting a diverse array of repair pathways, we envisage that MutSγ serves a similar purpose in the context of DSB repair during mammalian meiosis.

## MATERIALS AND METHODS

### Generation of Msh5^GA^ mice

The mouse Msh5 genomic locus was cloned from a P1 mouse ES cell genomic library (Genome Systems) (Edelmann et al., 1999). A 3.6 kb genomic *HindIII* fragment of mouse *Msh5* spanning exons 17-25 was inserted into *pBluescript SK* vector. Positive clones were identified by PCR. The G596A mutation and an analytic *BlpI* restriction site, were generated by site-directed mutagenesis in exon 19. A loxP flanked PGK hygromycin/neomycin cassette was inserted into the *MscI* site in intron 19. The targeting vector was linearized at the single *NotI* site and electroporated into WW6 ES cells. After selection in hygromycin, resistant colonies were isolated and screened by PCR. Positive clones were identified and injected into C57BL/6J blastocysts to produce chimeric animals. The PGK hygromycin/neomycin cassette was deleted by Cre-loxP-mediated recombination after mating of chimeric mice to *Zp3Cre* recombinase transgenic females (*C57BL/6J*). F1 offspring were genotyped and heterozygote animals were intercrossed to generate F2 homozygous mutant *Msh5^GA/GA^* mice and appropriate controls. Previously generated *Msh5^-/+^* mice were used for cross breeding studies to provide *Msh5^-/-^* null mice for comparison (Edelmann *et al.* 1999). All *Msh5^+/+^*, *Msh5^-/-^* and *Msh5^GA/GA^* mice used in these studies were backcrossed more than 10 times onto a *C57BL/6J* genetic background.

### Genotyping

Reverse transcription-PCR was performed on total RNA isolated from mouse tails with forward primer (5’-18d-3’) located in exon 18 and reverse primer 5’-TTGGTGGCTACAAAGACGTG-3’ located in exon 22 using the One Tube reverse transcription-PCR reaction kit (Roche) according to the manufacturer’s instructions. The following cycling conditions were used: 30 min at 50°C (1 cycle); 2 min at 94°C, 45 s at 60°C, and 45 s at 68°C (37 cycles); and 7 min at 68°C (1 cycle). The resulting 480 bp PCR product was subsequently restricted with *Blp*I.

### Care and use of experimental animals

Mice were maintained under strictly controlled conditions of light and temperature, with *ad libitum* access to food and water. All experiments were conducted with prior approval from the Cornell Institutional Animal Care and Use Committee. At least six mice per genotype were used for all studies.

### Histological analysis and TUNEL staining of mouse testis

Testes from 12 week-old mice were fixed in Bouin’s fixative for 6 hours at room temperature or 10% formalin overnight at 4°C, and then washed in 70% ethanol. Fixed and paraffin-embedded tissues were sectioned at 5 μm. Hematoxylin and eosin (H&E) staining and TUNEL staining and were performed as described previously (Holloway *et al.* 2008, 2010), the latter using Apoptag-peroxidase kit (Millipore).

### Chromosome preparation and spreads

The testes were decapsulated and incubated in hypotonic extraction buffer (HEB; 30 mM Tris, pH 8.2, 50 mM sucrose, 17 mM trisodium citrate dihydrate, 5 mM EDTA, 0.5 mM DTT, and 0.5 mM PMSF) for 1 hour on ice. About three to five millimeters length of seminiferous tubule was transferred into a drop of 20 μl hypotonic sucrose (100 mM, pH 8.2). After adding another drop of 20 μl of sucrose the tubule was macerated and the cell suspension was pipetted up and down for about 3-4 times. Remaining tubule fragments were removed from the cell suspension. Slides were coated with 1% paraformaldehyde containing 0.15% Triton X. 20 μl of the cell suspension were dispersed across the surface of one slide containing a layer of fixative. Slides were transferred to a humid chamber for 1-2 hours at room temperature and then allowed to air dry. Slides were washed three times for 3 min (0.4% Kodak Photo-Flo 200 in water) and air dried and stored at −80°C until use, not longer than 2 weeks.

### Immunofluorescence

The slides were washed in 0.4% Kodak Photo-Flo 200 in PBS and 0.1% Triton X-100 in PBS for 5 minutes each, blocked for 10 minutes in 10% antibody dilution buffer (ADB) in PBS (ADB: 3% bovine serum albumin, 0.05% Triton in 1 × PBS) followed by an overnight incubation in primary antibodies (at varying concentrations in ADB; Supplementary Table 1) at room temperature in a humid chamber. Slides were washed as described earlier and incubated for 1 h at 37°C in secondary fluorochrome conjugated antibodies in the dark. Primary and secondary antibodies used are listed in Supplementary table 1. All secondary antibodies were raised specifically against Fc fraction, Fab-fraction purified and conjugated to Alexafluor 594, 488, or 647.

### FIJI Image J Macro for SYCP1 & SYCP3 track measurements

TIFF images: Blue=DAPI, Red= SYCP3, Green=SYCP1

CZI images: (varies for every image batch) C1=Blue, C2=Red, C3=Green

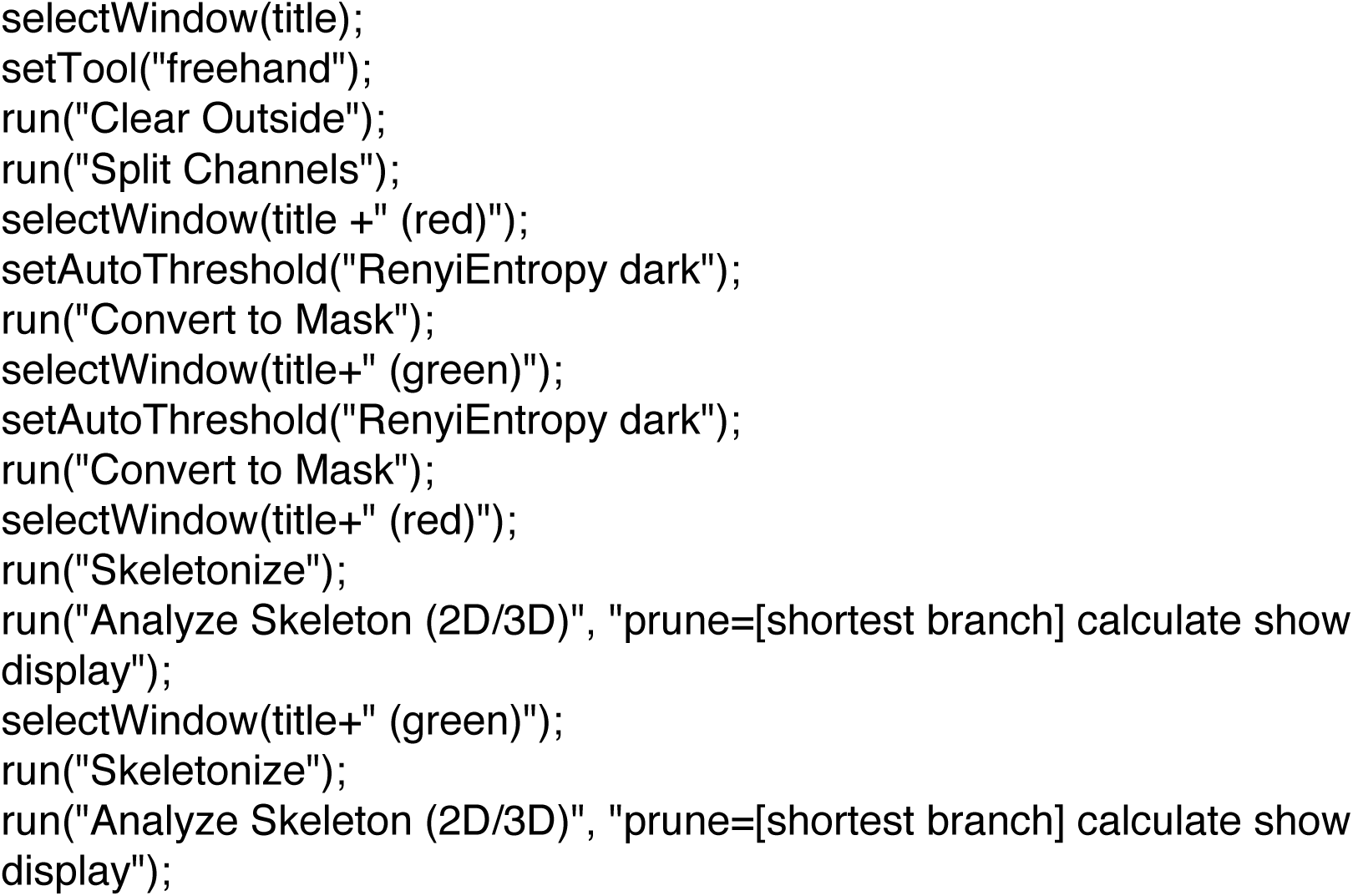

### Spermatocyte diakinesis spread preparations to observe chiasmata

Diakinesis chromosome spreads were prepared as previously described (Holloway *et al.* 2008, 2014). Slides were stained with 20% Giemsa for 2.5 min, washed, air-dried and mounted with Permount.

## ACKNOWLEDGEMENTS

The authors acknowledge, with extreme gratitude, the technical support of Mr. Peter L. Borst. We thank the members of the Cohen lab for their technical advice and for critical feedback on this manuscript and the studies outlined herein. We thank Mary Ann Handel (Jackson Laboratory, Bar Harbor, Maine) for providing the anti-H1t antibody. This work was supported by funding from NIH/NICHD to P.E.C. (HD041012) and from NIH/NCI (CA76329) and the Feinberg Family Foundation to W.E. Mice, antibodies, and plasmids are available upon request. The authors affirm that all data necessary for confirming the conclusions of the article are present within the article, figures, and tables.

---

**Supplemental Figure 1.**
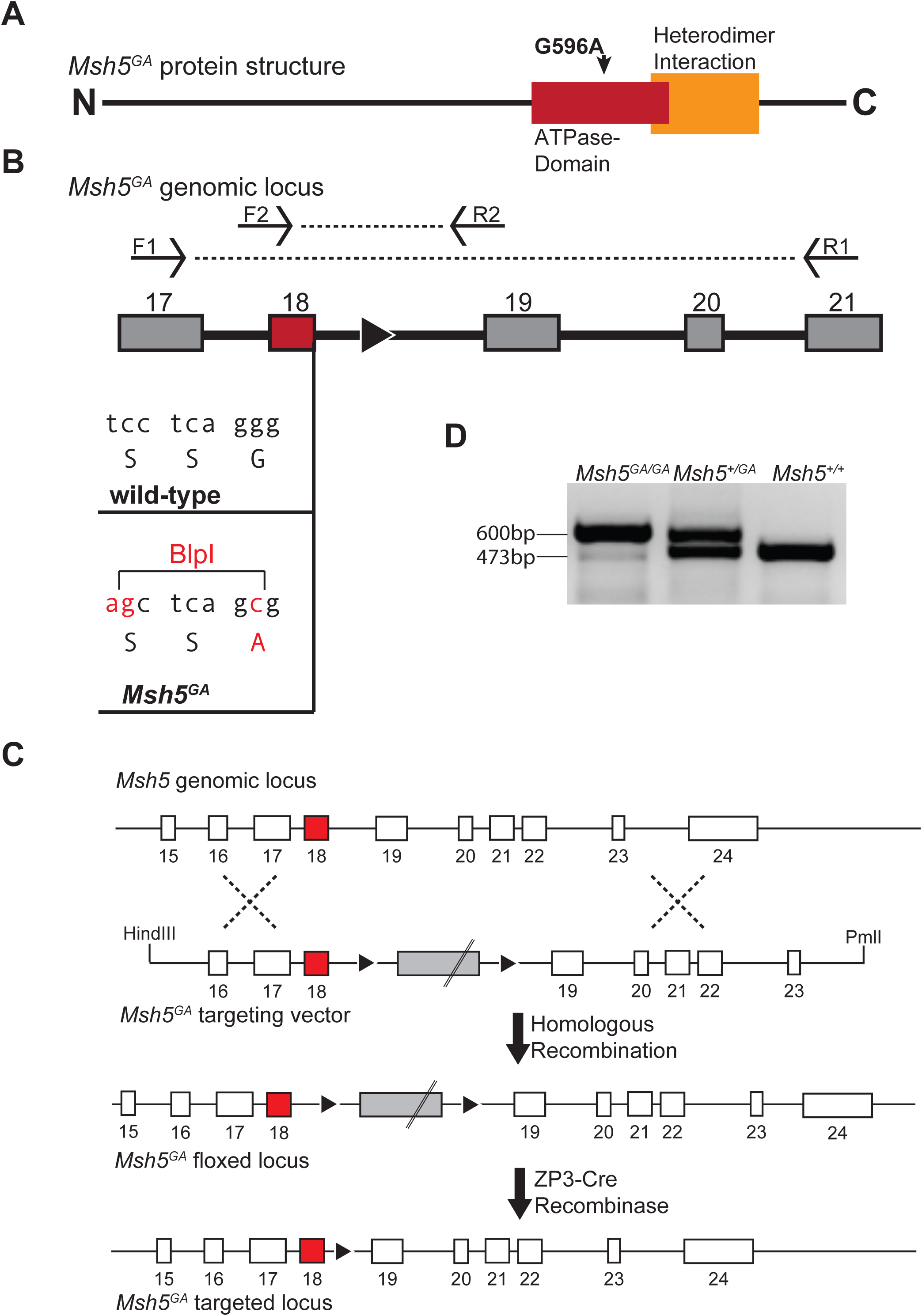
Description of the *Msh5^GA/GA^* mouse model. (a) Cartoon of *Msh5^GA^* predicted protein structure showing relative location of G596 point mutation in the ATPase domain. (b) Genotyping strategy for *Msh5^GA^* mice. Primers F1 [5’-CGGGACTACGGCTATTCGAGA-3’] and R1 [5’-GGCTACAAAGACGTGGGG-3’] are used for sequencing of the allele and F2 [5’-CAGGGTCAAAGTCATCACTG-3’] and R2 [5’-GGGCCATGAAAGTGATCAAG-3’] are used for genotyping. Novel RE site introduced in to exon 18 for additional genotyping conformation. (c) Gene targeting strategy used to introduce point mutation. (d) Example PCR results using primers F2 and R2 for *Msh5^+/+^*, *Msh5^+/GA^*, and *Msh5^GA/GA^* littermates.

**Supplemental Figure 2.**
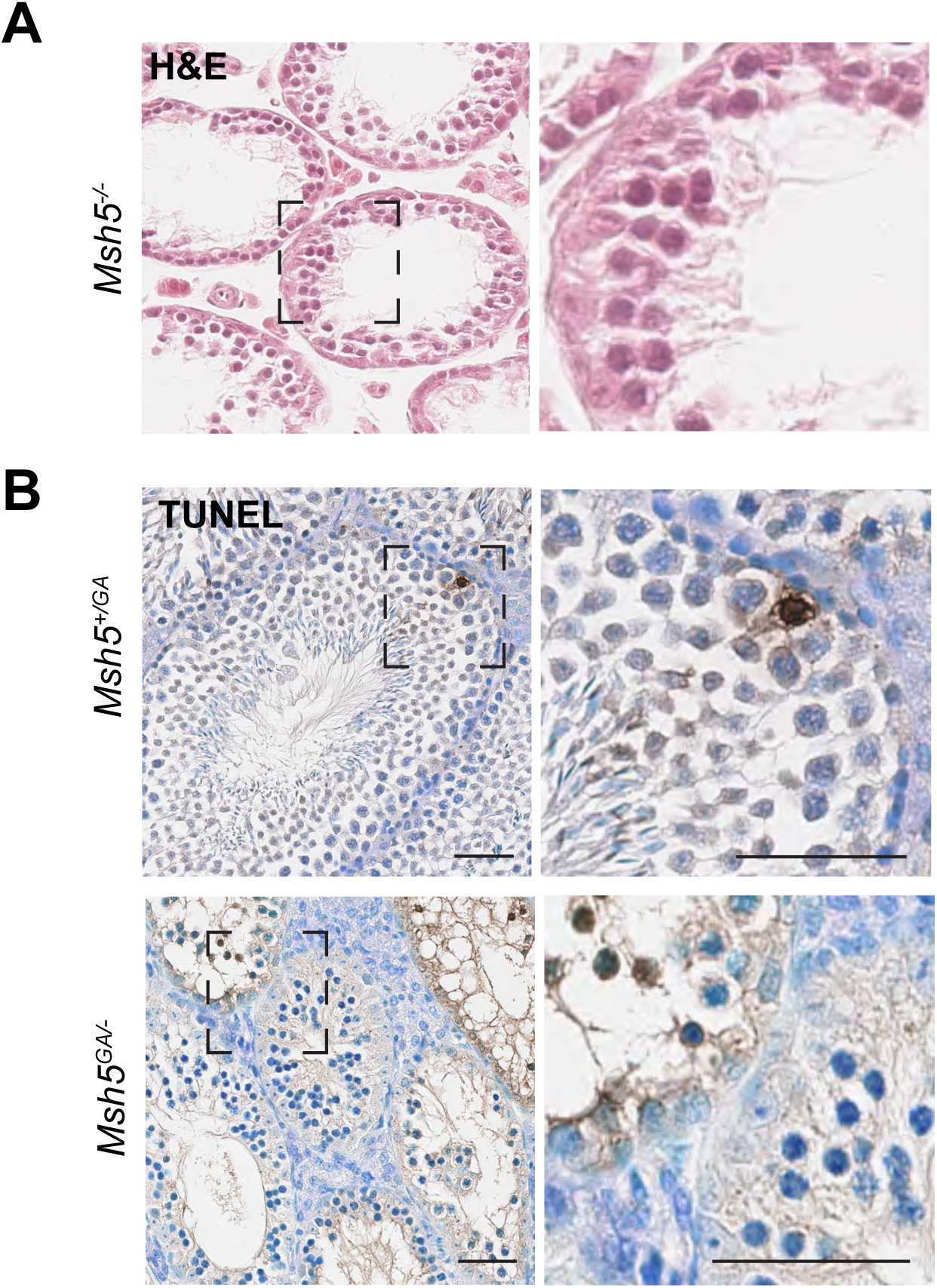
post-spermatocyte cell populations are absent in Msh5-/- histology and apoptotic cells are observed in *Msh5^GA/GA^* suggesting increased cell death. (a) H&E staining of adult paraffin embedded *Msh5^-/-^* testis sections lack cells beyond spermatocyte stage. (b) TUNEL of adult paraffin embedded *Msh5^+/+^*, *Msh5^+/GA^*, and *Msh5^GA/GA^* littermate testis sections fixed in 10% formaldehyde. Scale bars show 50 μM. Insets in right panels are show with increased magnification on the left panel.

**Supplemental Figure 3.**
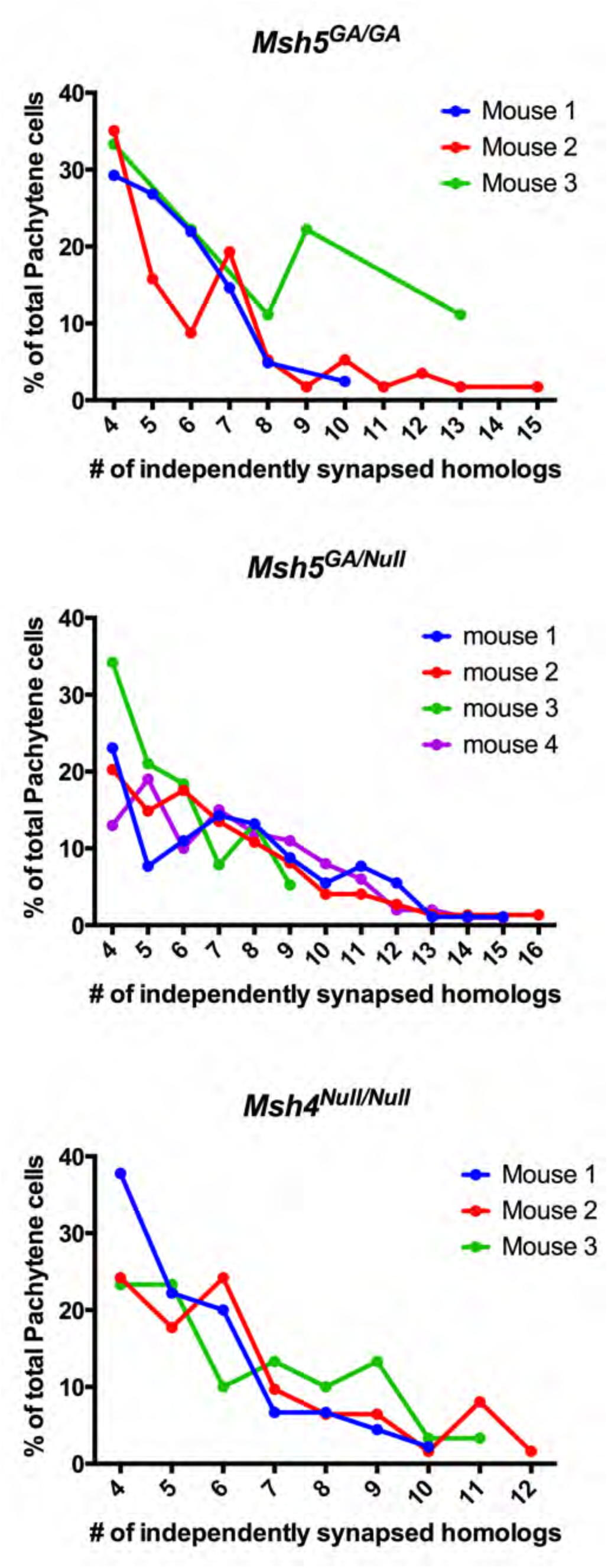
The frequency of independently synapsed homologs per pachytene-like cell in *Msh4^-/-^* and *Msh5^GA/GA^* males. The frequency of the number of independently synapsed homologs per pachytene-like cell for *Msh5^GA/GA^*, *Msh5^-/GA^*, and *Msh4^-/-^* in each of the three graphs respectively.

**Supplemental Figure 4.**
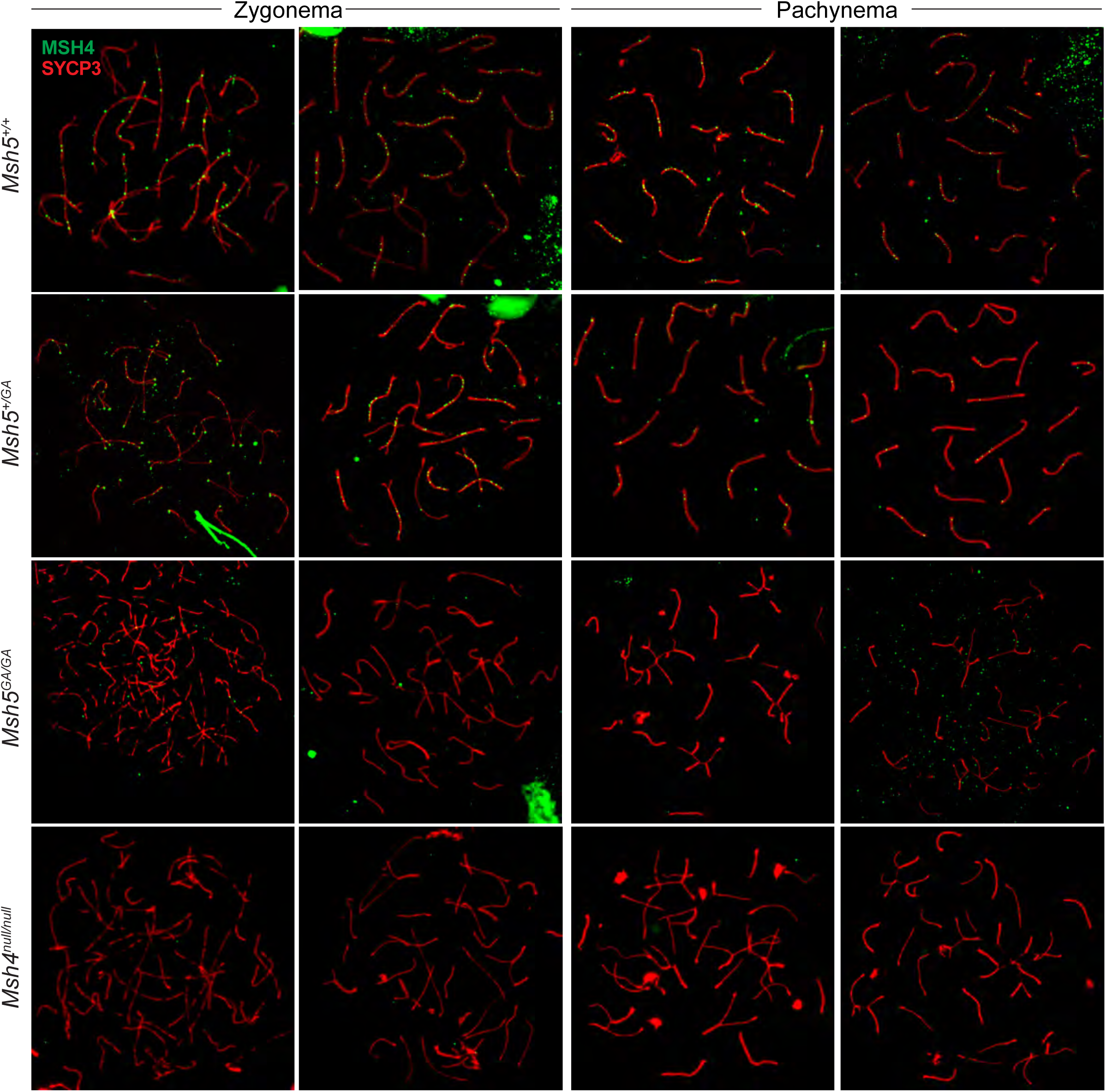
Additional exmaples of MSH4 localization in *Msh5^+/+^, Msh5^+/GA^, Msh5^GA/GA^, and Msh4^-/-^*. Localization of MSH4 (green) to SC (red) in adult *Msh5^+/+^*, *Msh5^+/GA^*, and *Msh5^GA/GA^* littermate chromosome spreads during zygonema and pachynema. Spreads from *Msh4^-/-^* animals are included for a negative control.

**Supplemental Figure 5.**
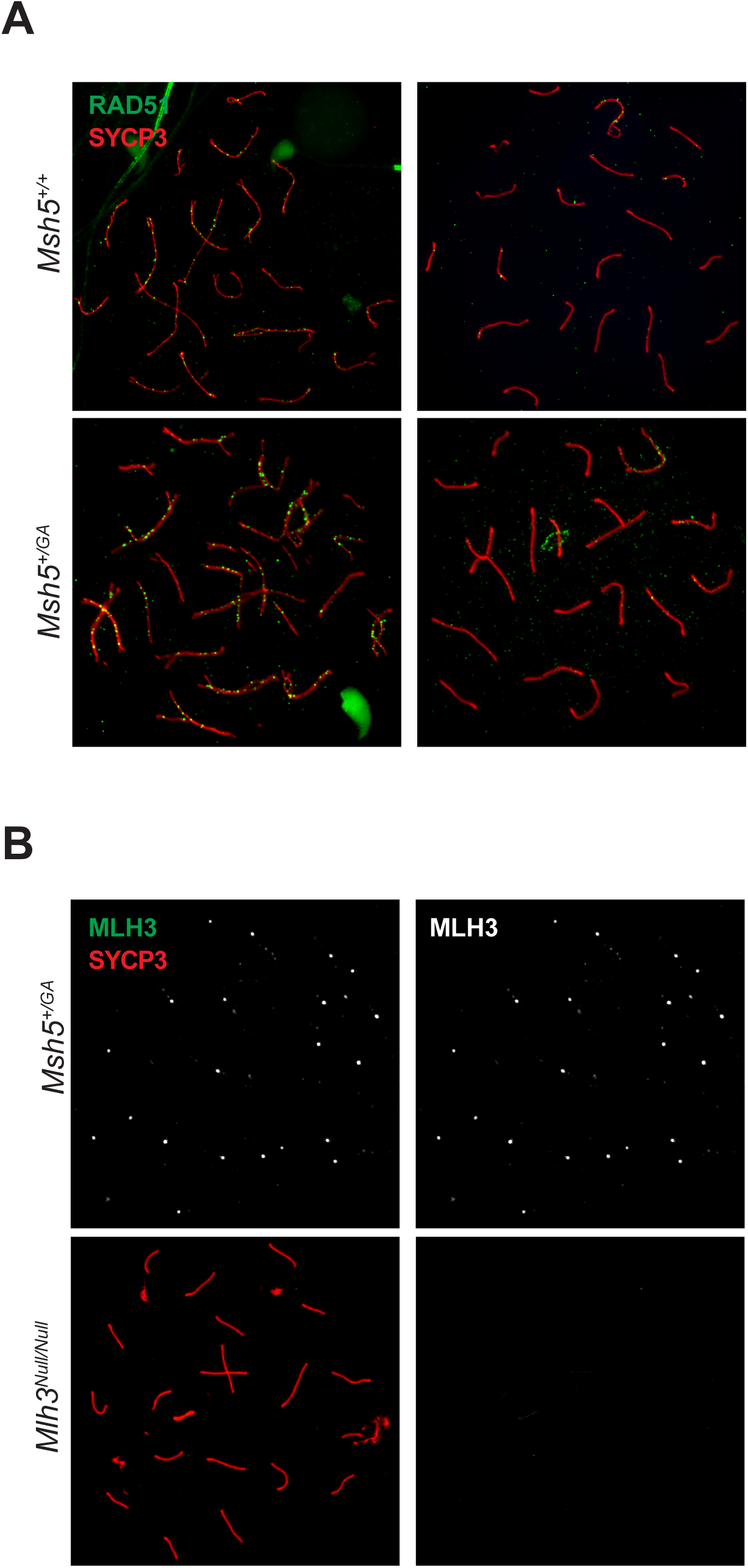
Heterozygous animals show normal RAD51 and MLH3 localization. (a) Immunofluorescent staining of DNA damage repair intermediate marker RAD51 (green) on synaptonemal complex proteins (red) in adult *Msh5^+/+^* and *Msh5^+/GA^* littermate chromosome spreads. (d) Immunofluorescent staining of MLH3 (green) in adult *Msh5^+/GA^* chromosome spreads show normal localization to SC (red) in pachytene, *Mlh3^-/-^* is included as a negative control.

